# The landscape of SARS-CoV-2 RNA modifications

**DOI:** 10.1101/2020.07.18.204362

**Authors:** Milad Miladi, Jonas Fuchs, Wolfgang Maier, Sebastian Weigang, Núria Díaz i Pedrosa, Lisa Weiss, Achim Lother, Anton Nekrutenko, Zsolt Ruzsics, Marcus Panning, Georg Kochs, Ralf Gilsbach, Björn Grüning

## Abstract

In 2019 the severe acute respiratory syndrome coronavirus 2 (SARS-CoV-2) caused the first documented cases of severe lung disease COVID-19. Since then, SARS-CoV-2 has been spreading around the globe resulting in a severe pandemic with over 500.000 fatalities and large economical and social disruptions in human societies. Gaining knowledge on how SARS-Cov-2 interacts with its host cells and causes COVID-19 is crucial for the intervention of novel therapeutic strategies. SARS-CoV-2, like other coronaviruses, is a positive-strand RNA virus. The viral RNA is modified by RNA-modifying enzymes provided by the host cell. Direct RNA sequencing (DRS) using nanopores enables unbiased sensing of canonical and modified RNA bases of the viral transcripts. In this work, we used DRS to precisely annotate the open reading frames and the landscape of SARS-CoV-2 RNA modifications. We provide the first DRS data of SARS-CoV-2 in infected human lung epithelial cells. From sequencing three isolates, we derive a robust identification of SARS-CoV-2 modification sites within a physiologically relevant host cell type. A comparison of our data with the DRS data from a previous SARS-CoV-2 isolate, both raised in monkey renal cells, reveals consistent RNA modifications across the viral genome. Conservation of the RNA modification pattern during progression of the current pandemic suggests that this pattern is likely essential for the life cycle of SARS-CoV-2 and represents a possible target for drug interventions.

## Introduction

Severe acute respiratory syndrome coronavirus 2 (SARS-CoV-2) is an RNA virus that causes coronavirus disease 2019 (COVID-19). The ongoing COVID-19 pandemic has put an enormous burden on human society in 2020 and is expected to have even longer-lasting impacts. Despite tremendous ongoing research efforts, we still do not have sufficient antiviral treatment solutions or a vaccine. Over the last two decades, the closely related zoonotic betacoronaviruses SARS-CoV and MERS have caused recurring outbreaks in the human population. The ability of coronaviruses (CoV) for cross-species transmission, their known reservoirs in multiple species, and their high replication rates keep CoVs a threat for the human population even beyond the 2020 pandemic. Understanding the molecular mechanisms behind the replication of SARS-CoV-2 is urgently needed.

SARS-CoV-2 carries an enveloped positive-sense single-stranded RNA genome (∼30kb) encoding a dense collection of structural and non-structural proteins (nsp), and accessory proteins. Like other members of the order Nidovirales, the genome encodes two polyproteins followed by a series of ORFs that are transcribed into sub-genomic RNAs (sgRNAs). Each transcribed sgRNA is thought to be translated into one protein, and its 3’ untranslated region overlaps with the coding sequence of the shorter downstream sgRNAs (1,2). Upon cell entry, ORF1a and ORF1b can be translated directly from the viral genome. A −1 ribosomal frameshifting upstream of the ORF1a stop codon allows the translation of ORF1b (3). The resulting polyproteins, pp1a and pp1b, are further cleaved by viral proteases and yield 11 and 15 nsps, respectively. The RNA-dependent RNA polymerase (RdRP) nsp12 performs the genome replication and the transcription of sgRNAs through negative-sense RNA template intermediates. To transcribe the sgRNAs, the negative RNA intermediates undergo discontinuous transcription, in which the RdRP skips the genome region between transcription-regulatory sequences (TRS) located at the 5’ end of the ORFs (TRS-B sites) and a corresponding TRS-Leader site at the 5’ end of the viral genome (for a review please see Sola *et al*. (1)). As a consequence, viral sgRNAs share a common 5’ leader sequence derived from the 5’ end of the genome up to the TRS-L site. Like host mRNAs, the viral genomic RNA and the sgRNAs have a methylated 5’ cap and a polyadenylated 3’ tail. Still, the transcriptomic aspect of CoVs, including SARS-CoV-2, is not fully understood. Transcript-level regulation of gene expression is widely used by the native cellular mechanisms of the host. Viruses have adopted and hijacked these mechanisms throughout their evolution (4). Understanding the biochemical characteristics of SARS-CoV-2 genomic and sgRNA molecules can provide valuable information for developing novel drug targets and optimizing the application of available therapeutics and mRNA-based vaccine development. The multifaceted functional aspects of RNA modifications have only recently been acknowledged and confirmed by several studies that have shown the important role of RNA modifications in the regulation of gene expression (5). Several studies indicate that RNA modifications play a pivotal role for viral infection and host defence (6,7).

More than 140 types of RNA modifications have been identified until now (8). While several protocols exist for the detection of nucleotide modifications such as RIP-seq, each assay can typically only identify one specific modification type. Raising specific antibodies to detect the growing number of known modifications remains an additional challenge (9). Direct RNA sequencing using Oxford Nanopore technologies (ONT) enables intermediate-free sensing of the nucleotides from the deviations in electrical signals while the RNA passes through the sequencing pores. The applicability of ONT-based solutions for detecting RNA modifications has been demonstrated in several studies (10), including work on SARS-CoV-1 (11).

Here, we study SARS-CoV-2 RNA modifications by direct sequencing of RNA from a human lung cell line infected with SARS-CoV-2. We present an extensive analysis of RNA modification patterns based on the sequencing of three virus isolates using two different modification prediction methods in a consistent manner. Furthermore, we reevaluate and compare our results to data from two previous reports of SARS-CoV-2 direct RNA sequencing experiments (12,13), which have analyzed the RNA modification of SARS-CoV-2 cultured in Vero (African green monkey kidney) cells, a cell line known to carry various chromosomal deletions and genetic rearrangements (14). Our analysis confirms and extends the previously reported results and, taken together, reveals that the transcripts of SARS-CoV-2 are consistently modified in different host cells.

## Results

### Cultivation of SARS-CoV-2 and RNA extraction

The aim of our study was to provide a replicate based direct RNA sequencing analysis of European SARS-CoV-2 to be able to analyze RNA modifications and predict the expressed viral transcripts. To this end, we cultivated SARS-CoV-2 isolates from three independent patients (*Fr1, Fr2, Fr3*) from Munich and Freiburg, including one of the first patients tested positive for SARS-CoV-2 in Germany. Isolate stocks obtained from infected Vero cell cultures were used to infect Calu-3 cells, a human lung epithelial cell line. We chose Calu-3 to study the viral RNA after infection of a disease-relevant human cell type. After 24 hours, the RNA of the infected cultures was extracted for deep sequencing. We applied classical short-read sequencing as well as direct RNA sequencing using nanopores. Short-read Illumina sequencing of the samples was essential to obtain a high-confidence list of genomic variants present in each isolate (Table S1). For nanopore sequencing, we used an ONT MinION sequencing device and sequenced poly-A enriched RNAs.

### Sequencing read statistics

The direct RNA sequencing experiments yielded a total of 2.3, 1.2 and 1.3 million sequencing reads for the three samples. We mapped the sequences of each dataset to the combination of the human host genome, the yeast enolase gene used as the ONT DRS spike, and the SARS-CoV-2 NCBI reference genome. Notably, between 62-70% of the mapped reads were mapped to the virus genome (Fig. 1a), which is very much comparable to the fraction of viral reads obtained by Kim *et al*. (12) using the Vero host cell line (Fig. S1a). In contrast to Calu-3 cells which were used for this study, Vero cells are interferon-deficient. Thus, our observation seems to indicate that the interferon deficiency of Vero cells does not benefit the viral life cycle to an experimentally relevant extent. This is in line with a recent study analyzing the host transcription response to SARS-CoV-2 infection (15).

**Figure 1:**
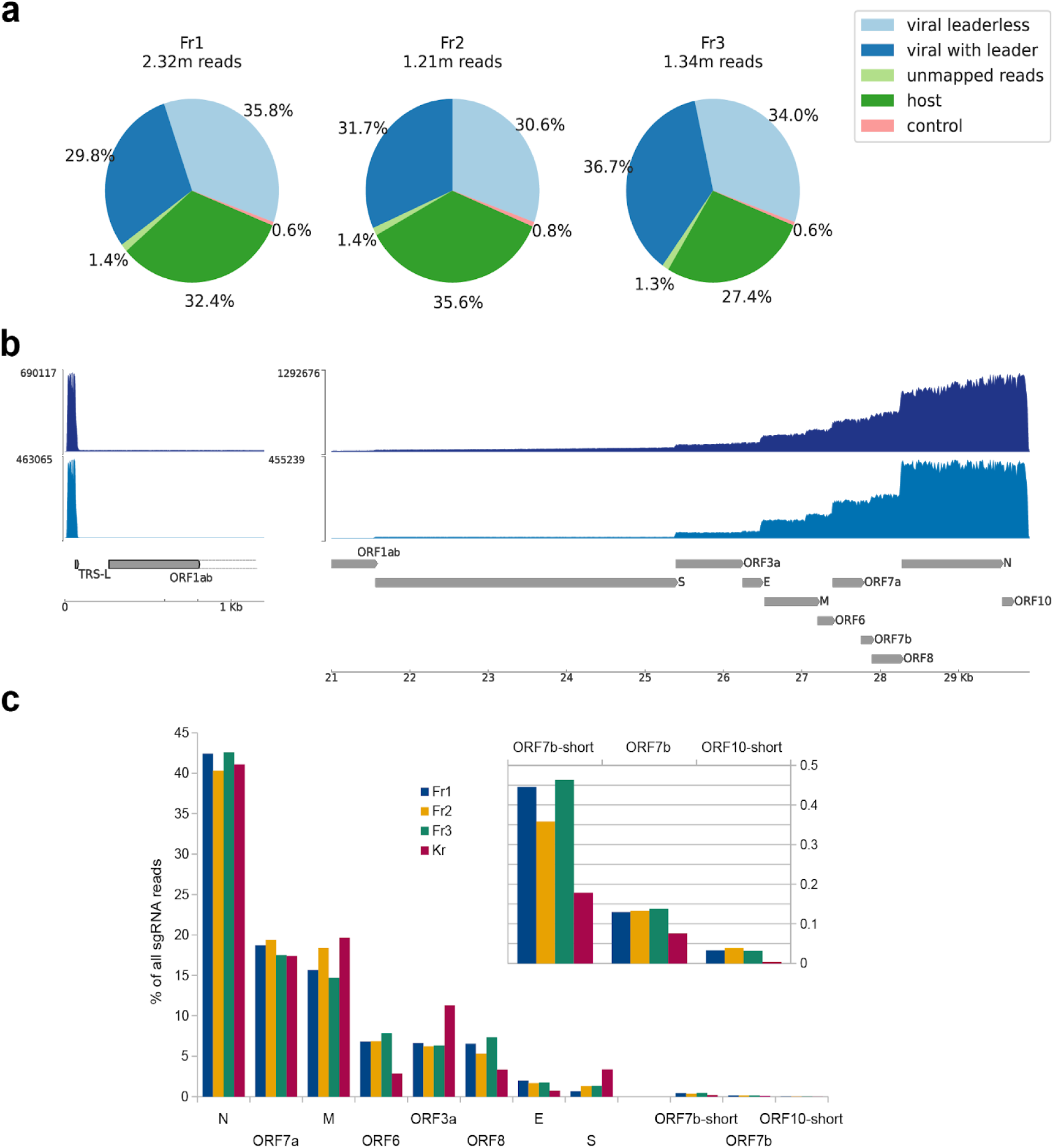
Direct RNA sequencing of SARS-CoV-2 infected Calu-3 human cell lines.**a:** Mapping statistics for reads obtained from human epithelial cells infected with three independent virus isolates. Given is the fraction of reads mapping to the human genome, ONT control ENO, and SARS-CoV-2. More than 60% of the mapped reads aligned to the virus genome. The subset of viral reads that span over the 5’ leader sequence are designated as *intact* reads. **b:** top panel, the coverage of viral reads across the SARS-CoV-2 genome with a truncated axis in case of ORF1ab. Bottom panel, the coverage of sgRNA reads with a leader sequence. **c:** Relative abundance of viral reads assigned to sgRNAs based on their support of canonical and newly observed TRS-B site usage. A linear scale is used to show the magnitude of expression differences. The inset shows a magnification of the three most lowly expressed sgRNAs. Fr1-3, three German virus isolates, this study; Kr, Korean isolate

### SARS-CoV-2 TRS-B sites and subgenomic RNAs

The long RNA sequencing reads generated for this study cover the entire SARS-CoV-2 genomic RNA as well as the different ORFs (Fig 1b,c, Fig. S1b). This allowed us to do an in-depth analysis of the genomic junctions, including the TRS-B sites described by Kim *et al*. (12). For comparability, we downloaded and reanalysed the DRS dataset published by Kim *et al*. and included it in our junction site analysis and downstream evaluations. Data from this dataset are designated as *Kr*.

Our scan of candidate landing regions upstream of predicted ORFs, and alternative start codons within them, monitored a total of 16 genomic regions and classified sequencing reads by the region they support (the result of this classification can be seen in Supplementary Table S2). Manual inspection of alignments of each class of reads enabled us to re-identify known TRS-B sites and to discover novel ones. We used this list of observed TRS-B sites for a more stringent classification of reads, which only considered reads supporting a junction between the TRS-L and one of the observed TRS-B sites. The results of this reclassification are shown in Table 1 and Figure 1c and confirm the existence of functionally active TRS-B sites upstream of all predicted ORFs except for ORF10. In agreement with Kim *et al*., we find evidence for an additional functionally active TRS-B site predicted to enable translation, from an alternative downstream start codon, of an ORF7b short isoform lacking the first 23 amino acids of the annotated protein.

**Table 1:** TRS sites for which evidence has been observed in this study. For each TRS we list the following: its position as 1-based start position of the core motif; its core sequence and the three bases flanking it on each side; the supporting read counts in each of the three samples from this study and in the reanalyzed Kr sample (for the TRS-L site, these counts are simply the sum of all the TRS-B counts since reads were required to support junctions between TRS-L and one of the TRS-B sites to be considered).

|  | Position | 5'-flank | core | 3'-flank | Fr1 | Fr2 | Fr3 | Kr |
| --- | --- | --- | --- | --- | --- | --- | --- | --- |
| TRS-L | 70 | TAA | ACGAAC | TTT | 638360 | 351025 | 445928 | 274213 |
| TRS-S | 21556 | TAA | ACGAAC | aaT | 4236 | 4571 | 5970 | 9202 |
| TRS-ORF3a | 25385 | TAA | ACGAAC | TTa | 42243 | 21827 | 28155 | 30943 |
| TRS-E | 26237 | agt | ACGAAC | TTa | 12572 | 5833 | 7741 | 2058 |
| TRS-M | 26473 | TAA | ACGAAC | Taa | 99958 | 64531 | 65542 | 53923 |
| TRS-ORF6 | 27040 | atc | ACGAAC | gcT | 43469 | 24048 | 34908 | 7866 |
| TRS-ORF7a | 27388 | TAA | ACGAAC | aac | 119472 | 68100 | 78032 | 47675 |
| TRS-ORF7b | 27674 | Ttc | AaGAAC | TTT | 829 | 467 | 616 | 208 |
| TRS-ORF7b-short | 27760 | TgA | ttGAAC | TTT | 2845 | 1258 | 2067 | 489 |
| TRS-ORF8 | 27888 | TAA | ACGAAC | aTg | 41675 | 18728 | 32794 | 9191 |
| TRS-N | 28259 | TAA | ACGAAC | aaa | 270852 | 141527 | 189961 | 112649 |
| TRS-ORF10-short | 29571 | TAA | ACG--- | TTT | 209 | 135 | 142 | 9 |

We compared the sequence contexts of the TRS-B sites of the two alternative ORF7b sgRNA isoforms with the TRS-L motif of the SARS-CoV-2 genome. We observed that transcription of sgRNAs for both the long and the short ORF7b isoform depends on imperfect TRS-B core motifs, AaGAAC and ttGAAC, respectively, instead of ACGAAC, which may explain the low number of observed reads supporting these two sgRNAs compared to those of other ORFs. In line with Kim *et al*., we do not find evidence for the predicted ORF10 sgRNA. However, we find weak support for an imperfect TRS-B site downstream of the presumed start codon of ORF10. The potential TRS-B site sequence, TAA ACG TTT carries a triplet deletion in the ACGAAC core motif, but shows identity to TRS-L in the three 5’ and 3’ flanking bases, respectively. Intriguingly, all three samples sequenced as part of this work have reasonable numbers of reads (> 100 per sample) supporting the usage of the site, while our reanalysis of the *Kr* sample confirmed a much smaller number (9) of such reads. sgRNA transcribed from this site could result in translation from an alternative start codon within ORF10 that would produce a peptide of just 18 amino acids compared to 38 of the predicted ORF10. Whether this potential peptide is of functional relevance remains to be validated.

### Detection of RNA modification

We mitigated the variability in the sgRNA expression levels and the ONT higher coverage bias at the 3’end of the transcripts by downsampling the collections of intact sgRNA reads. In this way, we get a quasi-uniform distribution of intact reads across all the samples and the sgRNAs except for ORF7b (Fig. S1b). For comparison, we also applied the same data processing workflows on datasets from Kim *et al*. and Taiaroa *et al*. (7,8).

We used the intact reads that were identified and down-sampled in the previous step for RNA modification detection by DRS using the two available in silico methods. For the identification of the modification sites, we used two different approaches for harnessing the sensed electrical signals from sequencing the native RNA molecules by nanopores. Typically, the electrical signal events aligned to positions, called squiggles, are compared between a condition with unknown putative modifications and a control condition. One strategy to detect the modified genomic positions is to compare the distribution of squiggles of two conditions, both encoding the transcripts of interest. Another strategy uses trained statistical models of the control condition to identify modification of the other condition by evaluating disagreement between the observed features and the model expectations.

Two sets of Galaxy workflows based on Tombo (16) and Nanocompore (17) tools were designed to compute the modification scores from the DRS data (Table S3). Both Tombo and Nanocompore support the distribution-based strategy while Tombo further can perform model-based modification detection. Since Nanocompore supports biological replicates, we used it as the distribution-based strategy for calling modifications from the three replicates (Fig. 2a). We further used Tombo to train models and calculate modification scores for individual samples (Fig. 2b). To this end, we utilized the in vitro transcribed (IVT) data of SARS-CoV-2 from Kim *et al*. as the unmodified RNA control dataset for both Tombo and Nanocompore. The distribution of the signals derived from virus RNA and unmodified RNA is representatively depicted for Fr3 in Figure 3.

**Figure 2:**
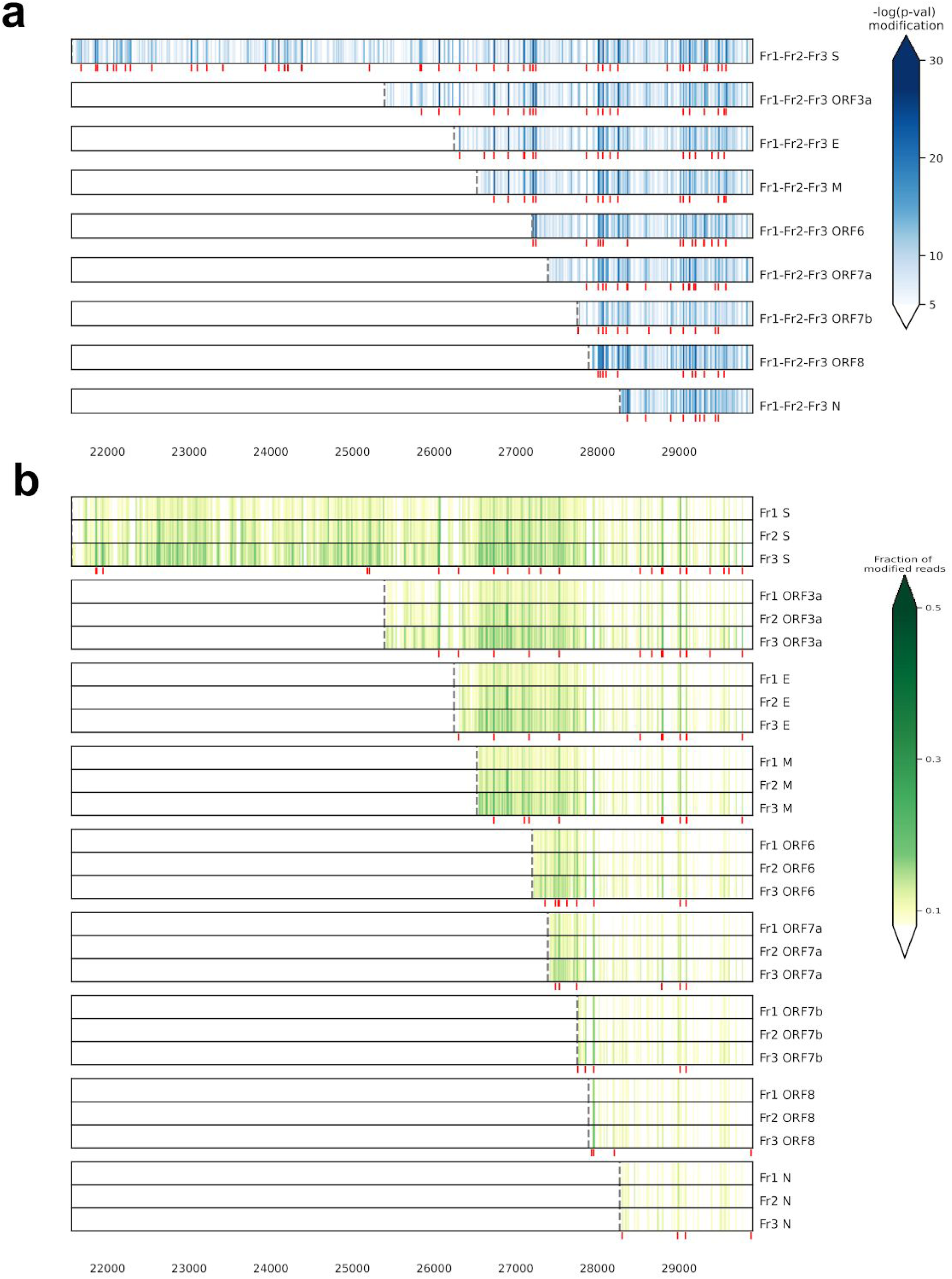
Detection of modified RNA bases in SARS-CoV-2 sgRNAs.**a:** Heatmaps of Nanocompore p-value scores for modified sites for the 3 sample replicates (Fr1-3) as compared to unmodified RNA data from Kim *et al*.. The genomic regions containing top-1% modification scores are marked in red. **b:** Heatmaps of the predicted fraction of modified bases using Tombo. The red marks show top-1% modified sites per sample that are common in at least two of the three samples.

**Figure 3:**
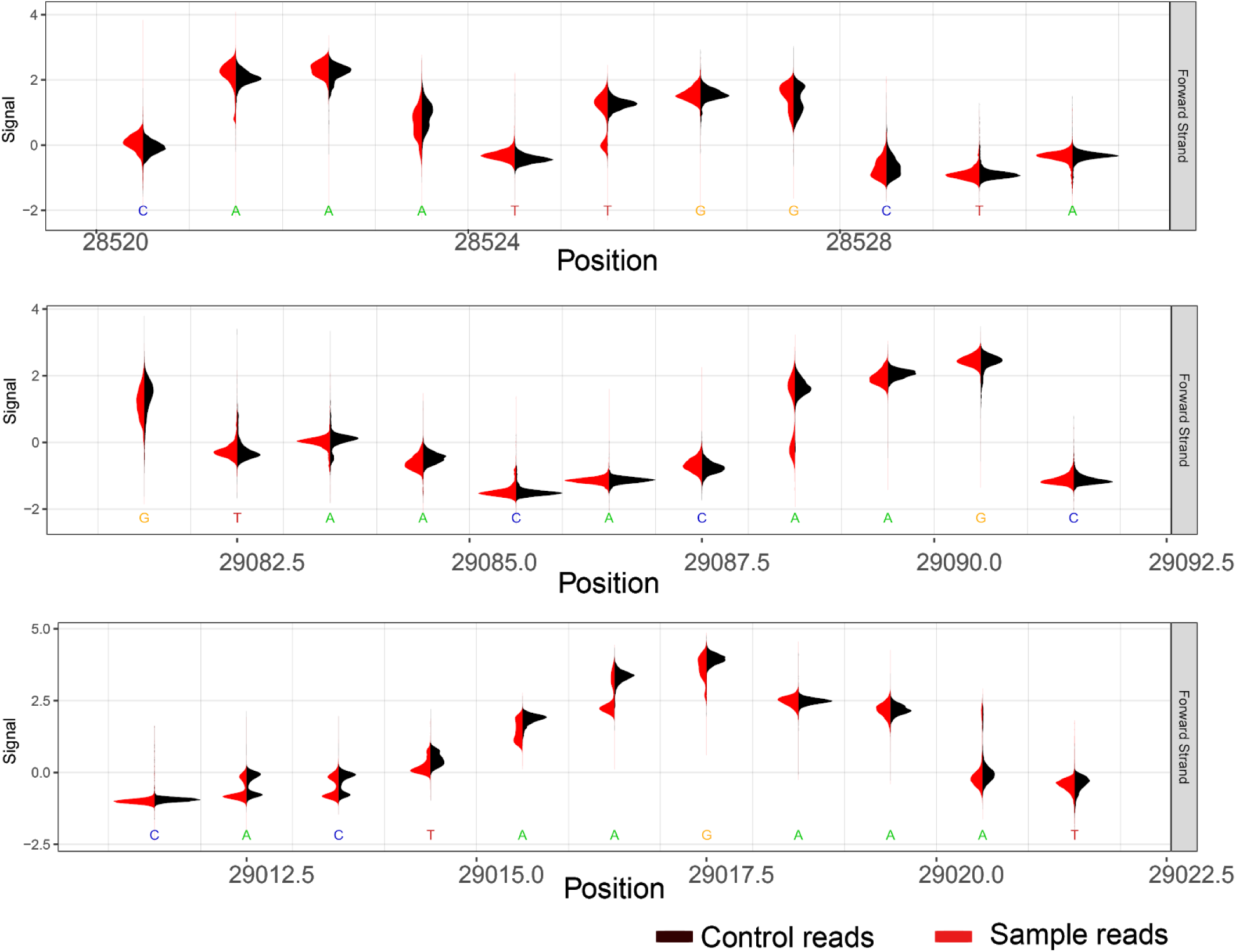
Distribution of nanopore measured ionic signals for exemplary regions with high modification scores according to Tombo and Nanocompore. Shown are signals obtained from unmodified RNA (black) and one representative sample, Fr3 (red).

### RNA modification sites of SARS-CoV-2 sgRNAs

We identified the positions modified in SARS-CoV-2 sgRNAs for all the sgRNAs and among all the datasets (Fig. 2a,b). We specifically focused on the modifications regions of the sub-genomic RNAs, i.e., the region downstream of the associated TRS-B sites. We excluded the genomic reads due to the moderately low number of intact reads. The modification results for 5’leader was also not considered due to the anomalies observed in the read coverage of the 5’leader site (Figure 1b).

By comparing the model-based prediction for the presented datasets (Fr1-3), we identified a high level of correlation between the modification rates of sgRNA positions in the three replicates (Figure 2b). This prompted us to perform a correlation analysis as depicted in Figure 4 representatively for sgRNA S and N. Notably, this analysis revealed a high correlation not only for the modification sites but also for the fractions of modification between biological replicates. We therefore tested the correlation between our data and the previously published data (Kr) (Fig. 4). Remarkably, the top-ranked modification sites are consistent and the correlation of the fraction of RNA modification fractions is high (Fig. 4), too. This observation was confirmed by visual inspection of raw signals (examples are shown in Fig. 5). We excluded the data of the Australian isolate from this analysis due to the relatively lower read coverage and different ratio of viral reads (Fig. S1a).

**Figure 4:**
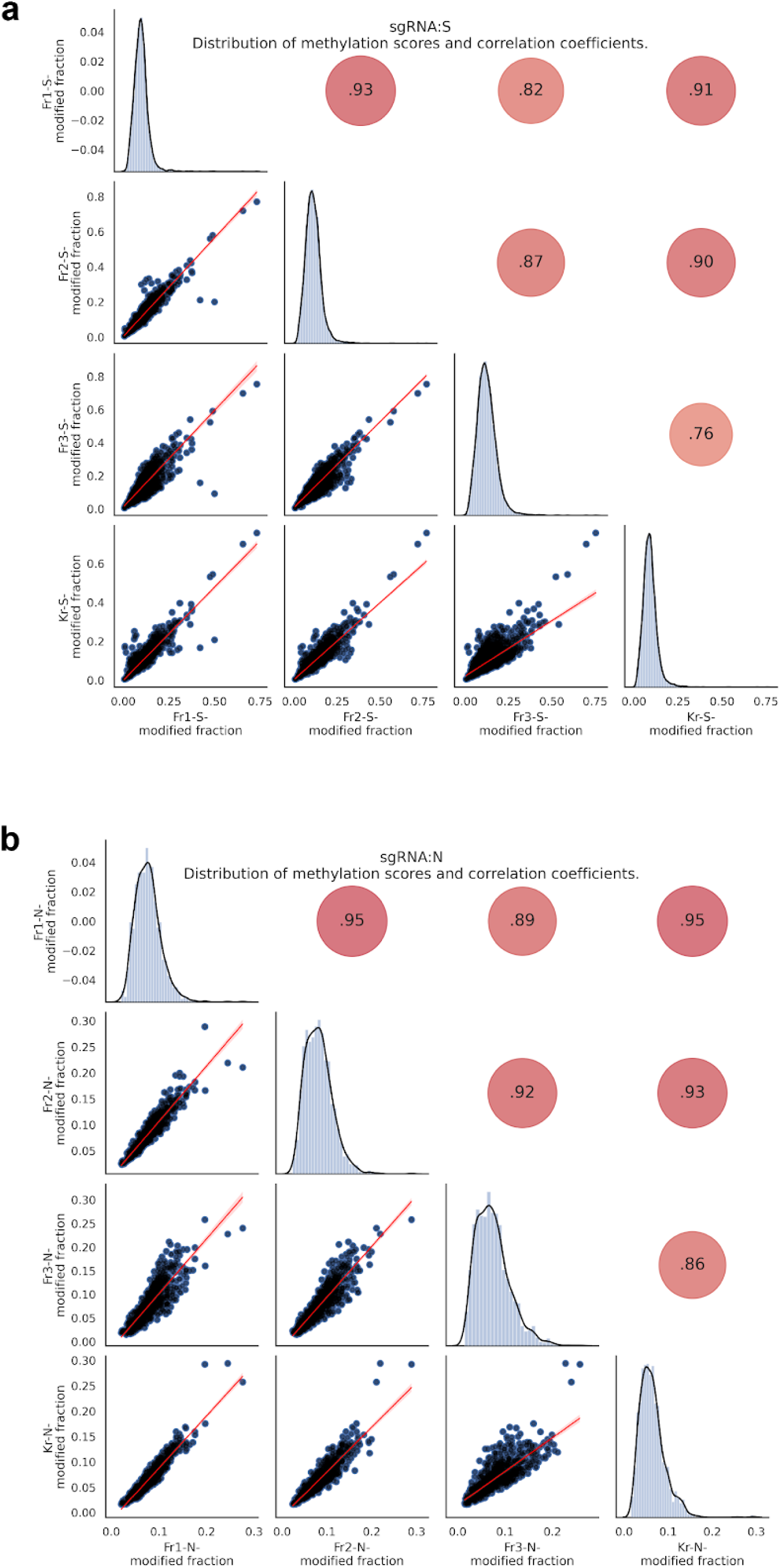
Correlation of the fraction of modified bases in the S **(a)** and N **(b)** sgRNAs computed using Tombo. Correlation coefficients are given in red circles.

**Figure 5:**
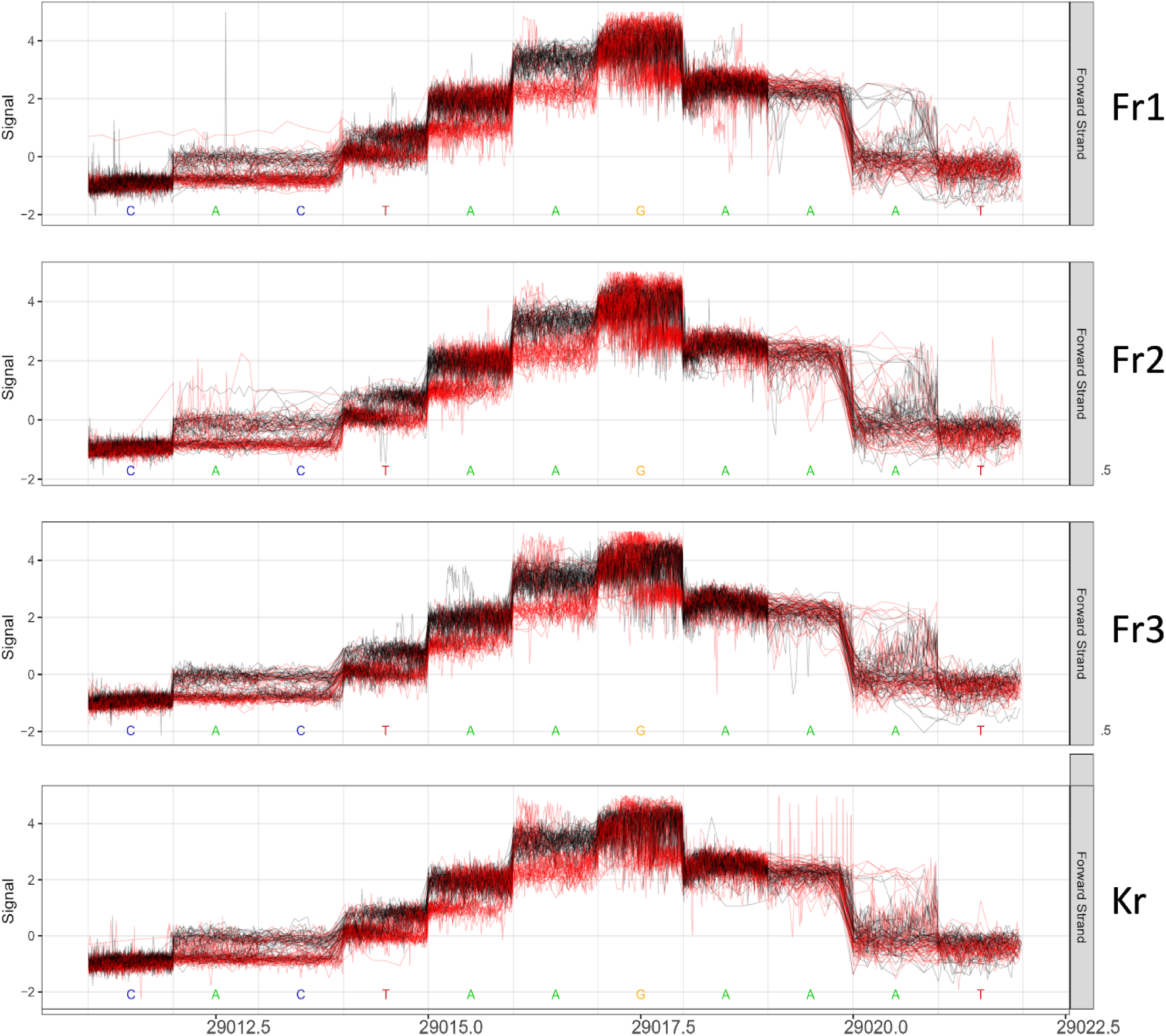
Direct RNA sequencing raw electrical signals of downsampled reads obtained from unmodified RNA (IVT, black), from samples generated for this study and from isolate from a published korean data set (Fr1-3 and Kr, red).

The large overlap of highly modified sites predicted by two independent algorithms supports the validity of our analysis and findings. However, for sites with a low modification ratio the predicted significance levels differ sometimes, indicating that additional biological replicates are needed to consistently reach a valid significance level.

## Conclusions

RNA modifications are essential modulators of RNA stability and function. The recent invention of direct RNA sequencing protocols using nanopores enable unbiased detection of RNA modification. In general, the analysis of DRS raw signals is challenging and not well standardized and thus only possible for experienced bioinformaticians. To enable more researchers to use this technology, we present two highly standardized analysis pipelines for DRS sequencing data. These pipelines were integrated into the Galaxy platform (18) and are accessible at https://covid19.galaxyproject.org/direct-rnaseq together with workflows for mapping reads to the viral genome, for calling genomic variants, and for identifying and extracting sgRNA-derived reads. Using these pipelines we analyzed the DRS data sets generated for this study, serving as the first DRS data from Europe, and compared it with the data from previous studies.

Here we generated the SARS-CoV-2 DRS sequencing data sets for the first time for three biological replicates. In contrast, to previously published data, viruses were cultured in a disease-relevant human epithelial lung cell line. Remarkably, the infection resulted in more than 60% of poly-A enriched RNA reads from SARS-CoV-2. We provide experimental evidence for transcription of a total of 11 sgRNAs, two of which are not part of the public SARS-CoV-2 reference annotation.

The comparative analysis of our three replicates with published data demonstrates a high degree of similarity between isolates from different continents and at both early and recent stages of the epidemie. Even the use of alternative host cells had little impact on the overall pattern of sgRNA transcription and RNA modifications. This high degree of conservation suggests that RNA modifications are relevant for the life cycle of SARS-CoV-2. Targeting of RNA modifying enzymes thus represents a novel therapeutic strategy. To test this hypothesis future studies have to identify and target the enzymes modifying the SARS-Cov-2 RNA and the associated RNA binding proteins. Although our results do not indicate the type of RNA modification, it provides a robust basis for detecting the different ribonucleoside modifications of SARS-CoV-2 in the future.

## Materials and methods

All work involving live SARS-CoV-2 was performed in the BSL-3 facility of the Institute of Virology, University Hospital Freiburg, and was approved according to the German Act of Genetic Engineering by the local authority (Regierungspräsidium Tübingen, permit UNI.FRK.05.16/05).

### Virus cultivation

SARS-CoV-2 isolates were propagated on VeroE6 cells (ATCC® CRL-1586) in Dulbecco’s Modified Eagle Medium (DMEM) with 2% FCS. For virus stocks, the cells were infected with a multiplication of infection (moi) of 0.001, supernatants were harvested after 50 h and aliquots stored at -80°C. Viral titers in the culture supernatants were determined using plaque-assays. The virus isolates used in this study were Muc-IMB-1/2020 (Bundeswehr Institute of Microbiology, Munich, Germany), FR/291.9/2020 and FR/291.13/2020 (Institute of Virology, Medical Center-University of Freiburg).

### Cells and infection

Calu-3 cells (ATCC® HTB-55™) were cultivated in DMEM supplemented with 5 % fetal bovine serum at 37°C and 5 % CO_2_. Cells were infected by washing confluent cells once with PBS and incubating them at a moi of 0.1 with virus preparations diluted in Opti-MEM for 1.5 h at RT. After the infection, fresh medium containing DMEM with 1 % FCS and 20 mM HEPES was supplied. The cells were harvested 24 h post infection to prepare total RNA.

### Viral RNA, total RNA and mRNA preparation

For Illumina cDNA RNA-seq, viral RNA was prepared from 200 µl of clarified virus stocks (3.000 rpm, 5 min) with the Quick-RNA Viral Kit (Zymo research) and eluted in 14 µl RNase free H_2_0. For Nanopore direct RNA sequencing, total RNA was isolated using the NucleoSpin RNA, Mini kit (macherey nagel) according to the manufacturers’ instructions. For each sample 1×10^6^ cells were lysed in 350 µl RA1 (supplemented with 3.5 μL ß-mercaptoethanol) and the RNA eluted in 50 µl RNase free H_2_0. RNA concentration and purity was quantified with a Qubit fluorometer (Quant-iT™ RNA HS Assay-Kit, ThermoFisher) and a Nanodrop (ThermoFisher), respectively. mRNA was prepared from total RNA by magnetic mRNA purification (Magnetic mRNA Isolation Kit,NEB) according to the manufacturer’s’ instructions with the following deviation: 50 µl purified total RNA (30-50 µg) was incubated with 450 µl Binding Buffer and added to 100 µl magnetic beads. The mRNA was eluted in 55 µl EB Buffer (Qiagen). To concentrate the mRNA, 99 µl Agencourt RNAclean XP (Beckman Coulter Life Sciences) were added and incubated at RT for 15 min. The magnetic beads were pelleted on a magnetic stand, washed twice in 70 % EtOH and dried for 5 - 10 min. Afterwards the mRNA was eluted in 11 µl RNase free H_2_0.

### Illumina cDNA RNA-seq

RNA-seq libraries (TruSeq® Stranded Total RNA Library Prep Human/Mouse/Rat, Illumina) were prepared from 150 ng of previously isolated viral RNA according to the manufacturers’ protocol. 10 pM pooled libraries were loaded onto a MiSeq cartridge (MiSeq Reagent Kit v2, Illumina) and run on a MiSeq (paired end, 300 cycles).

### Nanopore direct RNA sequencing

0,5 -1 µg of purified mRNA was subjected to direct RNA library preparation (SQK-RNA002, Oxford Nanopore Technologies) following the manufacturers’ instructions with the following deviations: Superscript IV (ThermoFisher) instead of Superscript III was used and the reverse transcription was performed for 2 h. The final library was loaded on a FLO-MIN106 flowcell and sequenced on a MinION (Oxford Nanopore Technologies) for 48 - 72 h, depending on the active channel count (MinKnow v3.6.5, Guppy v3.2.10).

### Quantification and Statistical analysis

#### Availability of analysis workflows and input data

The development of all analysis workflows used for the bioinformatic evaluation of the sequencing data was carried as part of the Covid-19 initiative of the Galaxy project (19). All Galaxy workflows and additional required inputs to them (beyond the sequencing data) are available from the Direct RNAseq subpage of the project at https://covid19.galaxyproject.org/direct-rnaseq.

#### Assignment of sequenced reads to viral transcripts

Mapping of the sequence reads to the corresponding genomes, extraction of *intact* reads and assignment to the sgRNAs were performed on the European Galaxy server.

For mapping, the ONT reads of each sample were first mapped to a virtual genome combined of the host (hg38) and the SARS-CoV-2 reference (NC_045512.2) genomes, as well as host rDNA (U13369.1) and ENO spike sequence using Minimap2 (20). The subset of reads that mapped to the viral genome was isolated using samtools (21) and served as input for a second round of mapping to only the viral genome and Minimap2 parameters optimized for the alignment of viral cross-junction sequences similar to Kim *et al*.. The complete mapping steps can be reproduced using our *Read mapping to viral genome* Galaxy workflow.

For the extraction of intact reads carrying the viral leader sequence and assignment of these reads to viral sgRNAs, we used a two-step strategy. First, we used bedtools (22) and samtools to extract reads, for which the mapping supported a junction between the viral TRS-L site and putative landing regions upstream of any potential longer ORF beyond ORF1ab. The list of landing region candidates used at this step includes the regions between each of the predicted structural ORFs and the next intervening upstream start codon, but also correspondingly defined regions upstream of potential alternative start codons within the S, 3a, M, 7b, N and 10 ORFs and enables a relatively unbiased detection of junction sites independent of prior assumptions about TRS-B sites.

Next, we inspected the resulting reads classifications with IGV (23) for evidence of junction events and used this information to build a list of TRS-B sites the use of which is supported by the sequencing data. This list was then used in a second round of assignment of reads to viral sgRNAs, in which only reads supporting a junction event between the TRS-L site and any of the confirmed TRS-B sites (with 10 flanking bases on each side to account for alignment ambiguities around the junction sites) were considered.

Both read classification strategies can be reproduced using Galaxy workflows to *classify ONT reads by candidate junction* and to *classify ONT reads by confirmed junction sites*, respectively. We have also made available the complete list of landing region candidates and confirmed TRS-B site regions used in these workflows.

#### Genomic variant analysis and masking of isolate variant sites

Genomic variants present in the viral isolates were identified from the MiSeq-sequenced reads data using the Galaxy workflow for variation analysis with paired-end data previously developed for the Covid-19 initiative of the Galaxy project. The exact version of the workflow used for the analyses described here is available together with the other workflows used in this study. A list of consensus variants identified in the three samples can be found as Supplementary Table S2.

Before computing RNA the modification score, the union of these identified variant sites for Fr1-3 plus the variants reported for Kr and Au samples were masked to avoid reporting mutations as false-positive modification signals. The genomic regions posing a high deviation in the coverage due to the overlaps at the boundaries of synthetic in vitro transcribed oligonucleotides in data from Kim *et al*. were further masked.

#### RNA modification detection

The collections of FASTQ-formatted intact reads with the viral leader sequence were used as input to Tombo. First, `tombo preprocess` and `tombo resquiggle` commands were invoked on the FASTQ files and the associated FAST5 collection (option --rna). Tombo detect_modification was invoked using the subcommand `model_sample_compare` (options --fishers-method-context 2 --minimum-test-reads 20 --sample-only-estimates) on the re-squiggled viral reads and the downsampled IVT data from Kim *et al*. The subcommand level_sample_compare was also applied with the same configuration (data not shown). The methylation scores were extracted from the computed statistics using the subcommand `text_output browser_files` --file-types dampened_fraction. The plots for ionic signals were also generated using Tombo.

The second workflow for distribution-based comparison of conditions was developed in Galaxy using Nanocompore and Nanopolish (17,24). To align the raw sequencing event data to the reference genome, Nanopolish subcommand `eventalign` was used (options --samples --scale-events --print-read-names) (17). The alignments produced in the previous step in BAM format and the associated reads in fastq format were provided to the Nanopolish tool. In the next step, the tabular output of event alignment was treated by removing the rows for the portion of the events that were aligned to the first 100 positions of the genome that covers the leader region using awk. This step has been necessary to have a proper utilization of Nanocompore tool that does not natively support spliced alignments. In the next step the event_align data was processed using NanopolishComp (https://github.com/a-slide/NanopolishComp) followed by Nanocompore `sampcomp` (options --sequence_context 2 --logit) to obtain the methylation scores. The p-value score GMM_logit_pvalue_context_2 was used to predict methylation.

## Acknowledgements

We are very thankful to the entire Galaxy community which is maintaining a vast collection of NGS-tools many of which have been fundamental for realizing this project. We thank the University Medical Center Freiburg for all the support and openness. We thank Hervé Menager supporting us when experiments were demanding more time and Nathan Roach and Stephan Flemming for extending our long-read-tool portfolio. We thank the Bundeswehr Institute of Microbiology, Munich, Germany, for providing us with SARS-CoV-2 isolate Muc-IMB-1/2020. We are very grateful to Oxford Nanopore Technologies for the awesome support and fast delivery - really impressive.

---

**Figure S1.**
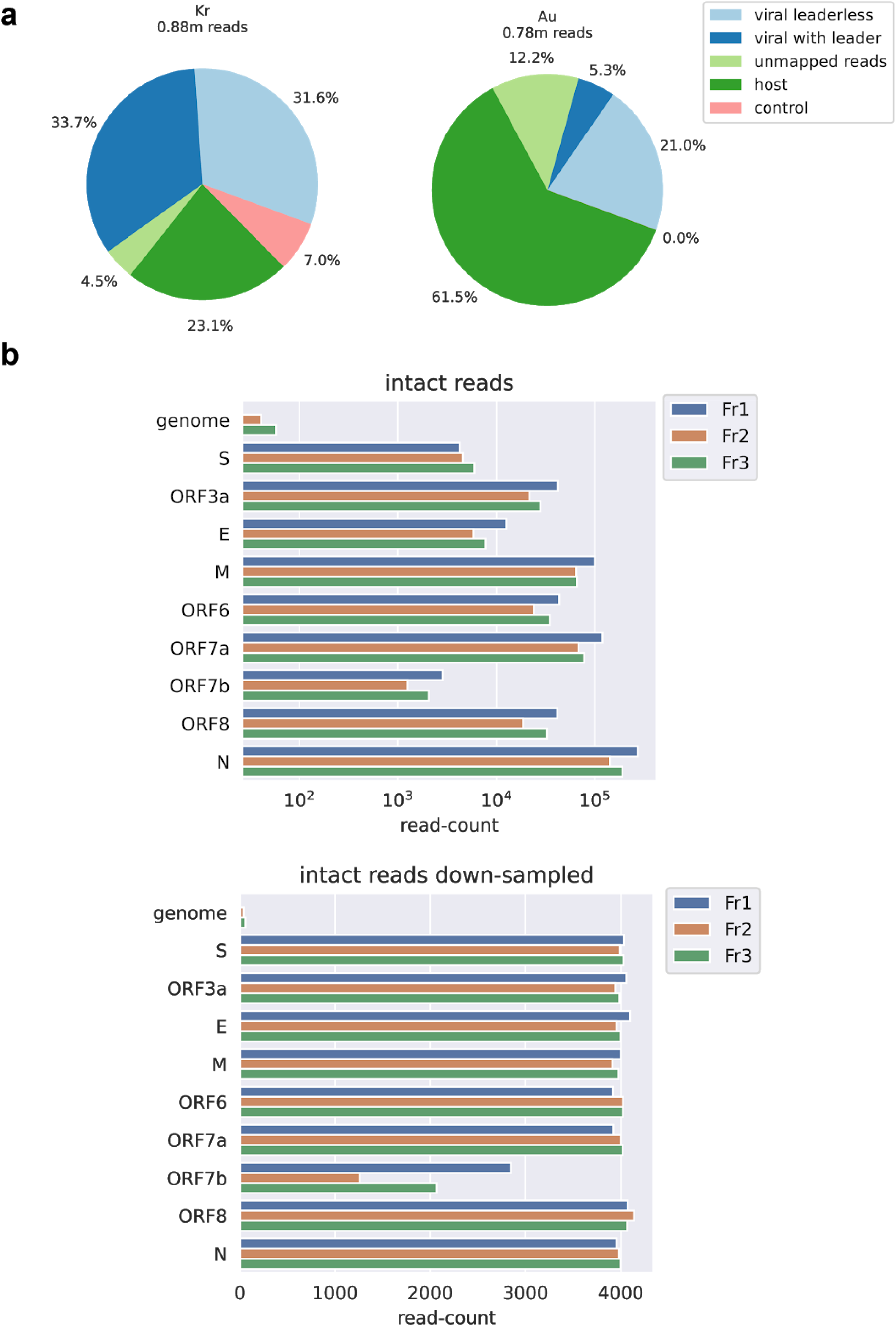
Mapping statistics of data sets and read distributions among the genomes.**a:** Mapping statistics of DRS reads for the human genome, ONT control ENO, and SARS-CoV-2. Depicted are results obtained for published data sets from Korea (Kr) and Australis (Au). **b:** Top panel, the total number of reads with a 5’leader sequence for the different sgRNAs and the genome. Bottom panel, the to maximal 4000 reads downsampled sgRNA reads for the downstream modification analysis.

## Supplementary Tables

**Table S1:**
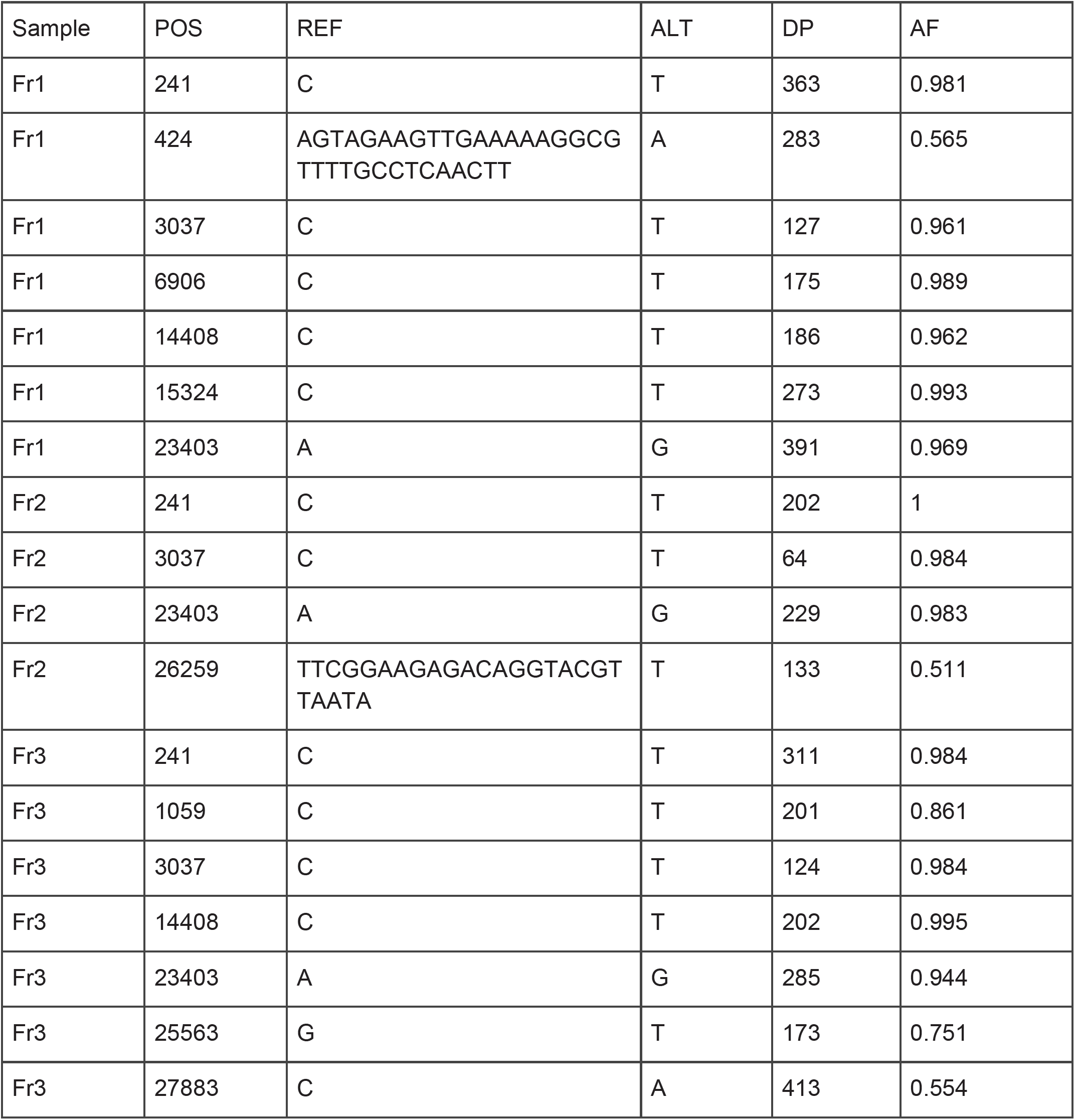
Genomic variants detected in the three studied isolates. To be included in this list the variant site had to show a depth of coverage (DP) > 10 and an alternate allele frequency (AF) > 0.5.

**Table S2:** Counts of reads supporting junctions between TRS-L and each of 16 TRS-B candidate regions. Asterisks mark candidate regions with unconvincingly low number of reads that were not considered for further analysis.

|  | Fr1 | Fr2 | Fr3 | Kr |
| --- | --- | --- | --- | --- |
| TRS-S | 3259 | 3423 | 3909 | 7436 |
| TRS-S-short* | 6 | 1 | 2 | 6 |
| TRS-ORF3a | 43037 | 22303 | 29003 | 31176 |
| TRS-ORF3a-short* | 21 | 11 | 18 | 21 |
| TRS-E | 11448 | 5304 | 6796 | 1958 |
| TRS-M | 105417 | 68621 | 69694 | 59037 |
| TRS-M-short* | 147 | 111 | 124 | 41 |
| TRS-ORF6 | 48774 | 27015 | 40328 | 8485 |
| TRS-ORF7a | 107595 | 60533 | 65452 | 44428 |
| TRS-ORF7b | 3576 | 2178 | 2882 | 1018 |
| TRS-ORF7b-short | 3563 | 1623 | 2610 | 633 |
| TRS-ORF8 | 34453 | 15224 | 24229 | 8119 |
| TRS-N | 279653 | 146443 | 199247 | 114874 |
| TRS-N-short* | 78 | 55 | 50 | 21 |
| TRS-ORF10* | 0 | 0 | 2 | 0 |
| TRS-ORF10-short | 280 | 177 | 183 | 18 |

**Table S3:**
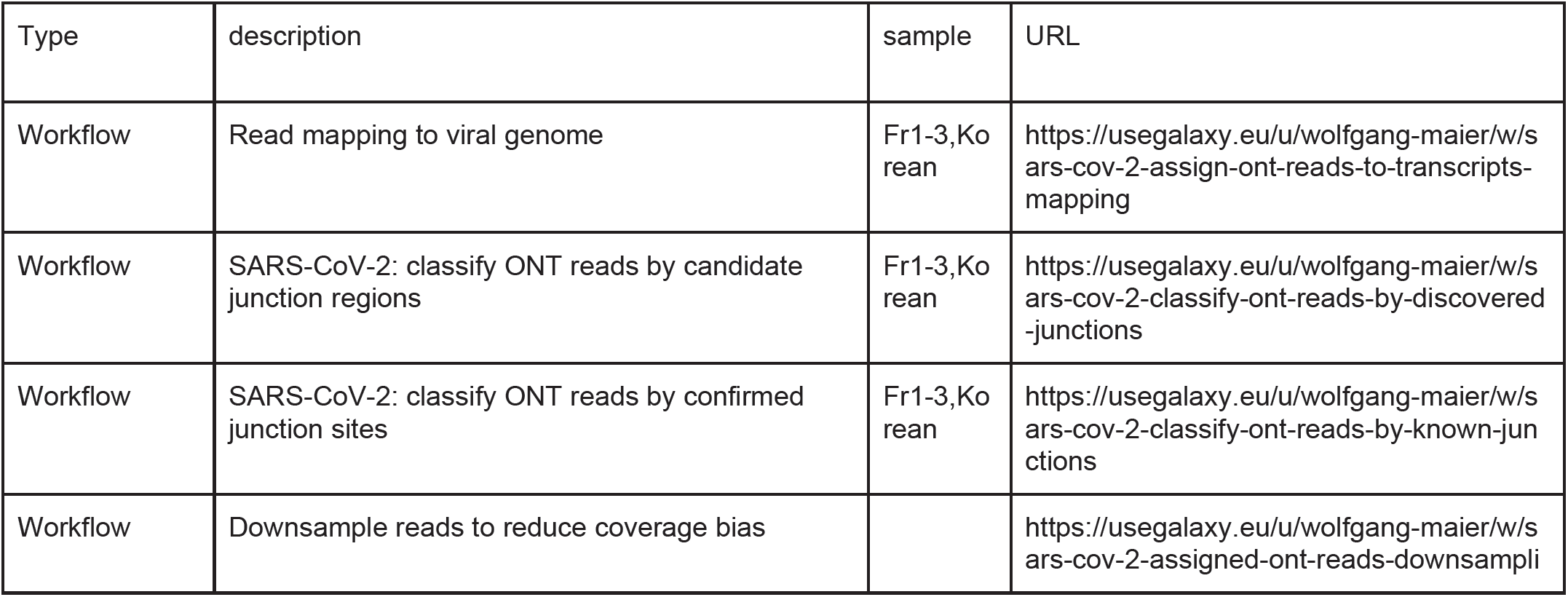
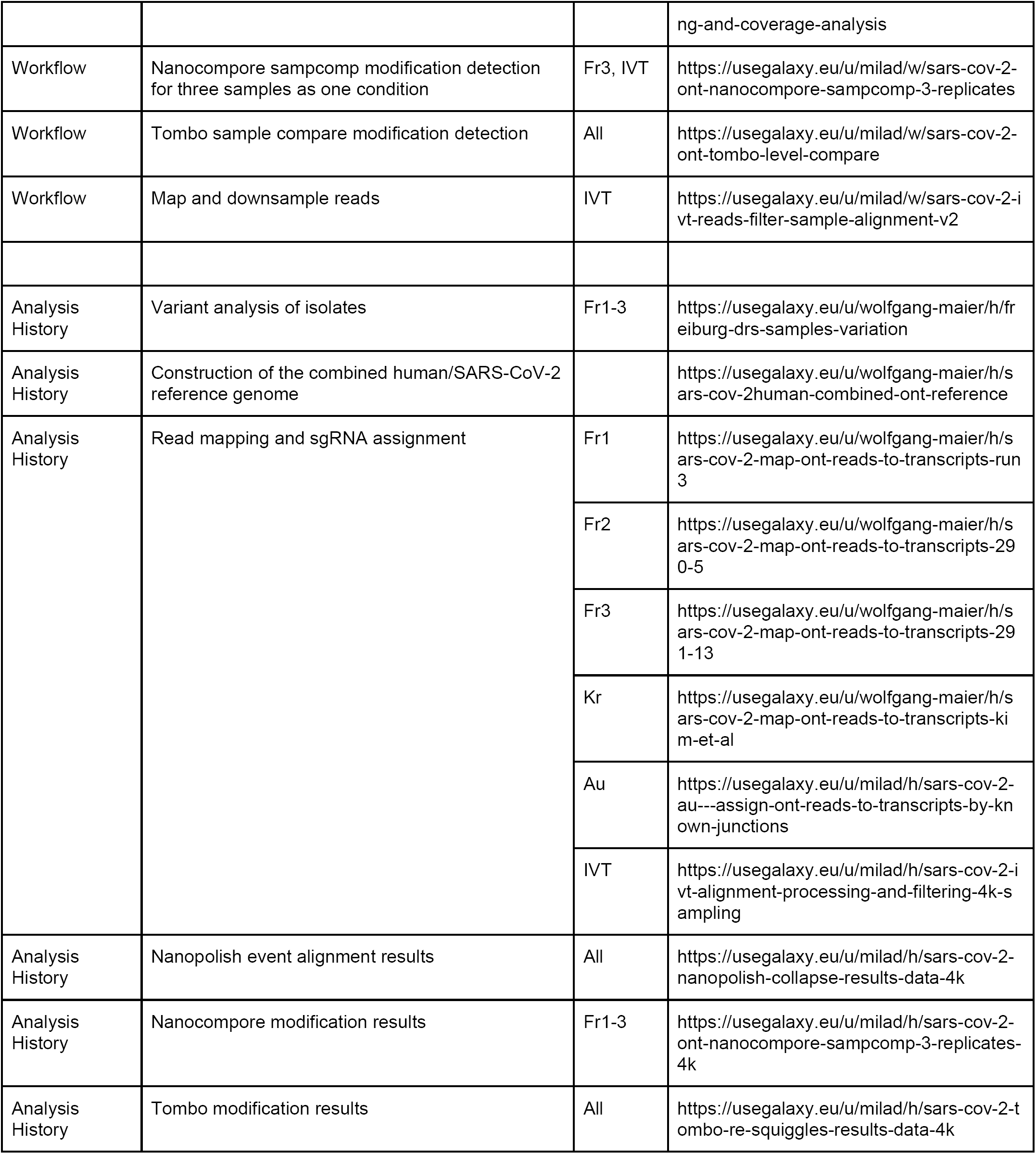
List of workflows and Galaxy histories containing all the work described in this study.

